# Identifying Priority Landscapes for Conservation of Snow Leopards in Pakistan

**DOI:** 10.1101/2020.01.27.920975

**Authors:** Shoaib Hameed, Jaffar ud Din, Hussain Ali, Muhammad Kabir, Muhammad Younas, Wang Hao, Richard Bischof, Muhammad Ali Nawaz

**Affiliations:** Carnivore Conservation Lab, Department of Animal Sciences, Quaid-i-Azam University, Islamabad, Pakistan; Snow Leopard Trust, Pakistan Program, Islamabad; Institute of Biological Sciences, Faculty of Science, University of Malaya, Kuala Lumpur, Malaysia; School of Life Sciences, Peking University, Beijing 100871, China; Faculty of Environmental Sciences and Natural Resource Management, Norwegian University of Life Sciences, Ås, Norway

**Keywords:** snow leopard, distribution, habitat, movement corridor, maxent, circuitscape, model landscape.

## Abstract

Pakistan’s total estimated snow leopard habitat is about 80,000 km^2^ of which about half is considered prime. However, this preliminary demarcation was not always in close agreement with the actual distribution—the discrepancy may be huge at the local and regional level. Recent technological developments like camera trapping and molecular genetics allow for collecting reliable presence records that could be used to construct realistic species distribution based on empirical data and advanced mathematical approaches like MaxEnt. Current study followed this approach to construct accurate distribution of the species in Pakistan. Moreover, movement corridors, among different landscapes, were also identified through the circuit theory. The habitat suitability map, generated from 384 presence points and 28 environmental variables, scored the snow leopard’s assumed range in Pakistan, from 0 to 0.97. A large shear of previously known range represented low-quality habitat, including areas in lower Chitral, Swat, Astore and Kashmir. Conversely, Khunjerab, Misgar, Chapursan, Qurumber, Broghil, and Central Karakoram represented high-quality habitats. Variables with higher contribution in the MaxEnt model were precipitation of driest month (34%), annual mean temperature (19.5%), mean diurnal range of temperature (9.8%), annual precipitation (9.4%) and river density (9.2). The validation texts suggest a good model fit, and strong prediction power.

The connectivity analysis revealed that the population in the Hindukush landscape appears to be more connected with the population in Afghanistan as compared to other populations in Pakistan. Similarly, the Pamir-Karakoram population is better connected with China and Tajikistan, while the Himalayan population was with the population in India.

Current study allows for proposing three model landscapes to be considered under GSLEP agenda as regional priority areas, to safeguard safeguard future of the species in the long run. These landsacpes fall in mountain ranges of the Himalaya, Hindu Kush and Karakoram-Pamir, respectively. We also identified gaps in existing protected areas network, and suggest new protected areas in Chitral and Gilgit-Baltistan to protect critical habitats of snow leopard in Pakistan.

## Introduction

The snow leopard, *Panthera uncia* has obtained an iconic status worldwide and is treated as a flagship species of the vast ecosystem of the Greater Himalayas [1]. The species is native to the mountain ranges of Central and Southern Asia—some of the world’s most rugged landscapes [2]. It occurs in the Hindu Kush, Karakoram, Altai, Sayan, Tien Shan, Kunlun, Pamir, and outer Himalayan ranges, and smaller isolated mountains in the Gobi region [3, 4]. Global range sizes vary from 1.2 million to over 3 million km^2^ [5] and the species is highly threatened throughout its range. A recent study estimated its occupied range to be about 2.8 million km² [6], across. 12 countries—Afghanistan, Bhutan, China, India, Kazakhstan, Kyrgyzstan, Mongolia, Nepal, Pakistan, Russia, Tajikistan and Uzbekistan ([7–9]. Potential range of snow leopard may also occurs in northern Myanmar, but recent snow leopard presence has not been confirmed [5].

Considering Roberts’ [10] range maps as a base, Pakistan’s total estimated snow leopard habitat is about 80,000 km^2^ of which about half is considered prime [11]. However, this preliminary evaluation of the snow leopard’s distribution is based on expert judgements, anecdotal information and topographic elements like terrain. Consequently, these distribution maps were not always in close agreement with actual distribution—the discrepancy may be huge at the regional and global level [7]. Accurate modelling of the geographic distribution of species is crucial to various applications in ecology and conservation [12, 13]. Conservationists often need precise assessments of species’ ranges and current species distribution patterns. Simple range description is essential, but identification of those factors that restrict distributions is also critical to promote the benefits of conservation management [14].

Factors that affect species distributions and habitat selection have great significance to researchers and wildlife managers. It is important to be aware of the influence of variables on species’ occurrence [15]. There is also a prepared source of environmental information, including global databases of climate and digital elevation models and species distribution models (SDMs), being used in ecological research and conservation planning [16]. Currently, ecological niche models (ENMs) and SDMs are increasingly being used to map potential distributions of many species [13]. These models incorporate species occurrence data with climatic and other environmental variables to produce reliable distribution maps of species [17] that are used to design scientific surveys and plan sustaible conservation [18]. The task of a modelling method is to predict the environmental suitability for the species as a function of given environmental variables [12].

Many models like BIOCLIM, BLOMAPPER, DIVA, DOMAIN, CLIMEX, GAM, GLM and GARP have been used in species distribution modelling [19, 20], but maximum entropy (MaxEnt) is argued to possess the best predictive capacity [21–23] and produces the most accurate distribution functions [23]. Several studies indicate that MaxEnt modelling performs better than other models [19]. MaxEnt estimates the probability of the presence of a species based on occurrence records and randomly generates background points by finding the maximum entropy distribution [18, 24].

These models can use either presence/absence data or presence-only data. The use of presence/absence data in wildlife management and biological surveys is widespread [25]. By contrast, absence data is often unavailable and difficult to verify given the potential for a species to be present at a site but not observed [26]. However, SDMs trained on presence-only data are frequently used in ecological research and conservation planning [16]. Understanding how predictions from presence/absence models relate to predictions from presence-only models is important because presence data is more reliable than absence data [27]. Presence-only modelling methods only require a set of known occurrences together with predictor variables such as topographic, climatic, and biogeographic variables [12].

Connectivity among habitats and populations is another critical factor that influences variety of ecological phenomena, including gene flow, metapopulation dynamics, demographic rescue, seed dispersal, infectious disease spread, range expansion, exotic invasion, population persistence and maintenance of biodiversity [28, 29]. Preserving and restoring connectivity is one of the top conservation priorities and conservation organizations are devoting substantial resources to accomplish these goals [30, 31]. A reliable, efficient and process-based approached is required to achieve this objective in complex landscapes. A new class of ecological connectivity models based on electrical circuit theory were introduced by McRae et al. [32]. Resistance, current and voltage calculated across graphs or raster grids can be associated to ecological processes like; individual movement and gene flow, that take place across large population networks or landscapes.

Given the multitude of threats to snow leopards and their habitat, it is imperative that comprehensive landscape-level conservation strategies be developed that are based on reliable information on species survival requirements. A global strategy to safeguard snow leopards and the vast ecosystem they inhabit—which includes 12 nations and supports 1 billion people—has already been established: The Global Snow Leopard Ecosystem Protection Program (GSLEP). Its overall aim is to secure at least 20 snow leopard model landscapes across the cat’s range by 2020 [33]. Under the GSLEP initiative, the selection of model landscapes requires a clear understanding of areas that represent the species’ prime habitat so that conservation efforts in the next decade can focus on securing areas that hold or have potential to hold larger populations. Recent technological developments like camera trapping and molecular genetics allow for collective reliable presence records that could be used to construct realistic species distribution based on empirical data and advanced mathematical approaches like MaxEnt. This study aims to support the GSLEP by identifying core habitats and movement corridors through upgrading knowledge on snow leopard distribution.

## Materials and Methods

### Study Area

The study focused on known snow leopard range in Pakistan which encompasses four high mountainous ranges; the Himalaya, Karakoram, Pamir and Hindu Kush spread across three administrative units, i.e. Khyber Pakhtunkhwa (KP), Gilgit-Baltistan (GB) and Azad Jammu and Kashmir (AJK). Targeting major protected areas and other potentially suitable habitats, we surveyed 20 sites with a collective area of around 40,000 km^2^ (**Fig 1**). The surveyed areas constitute 50% of reported snow leopard habitat in Pakistan (80,000 sq. km) [34].

**Fig 1.**
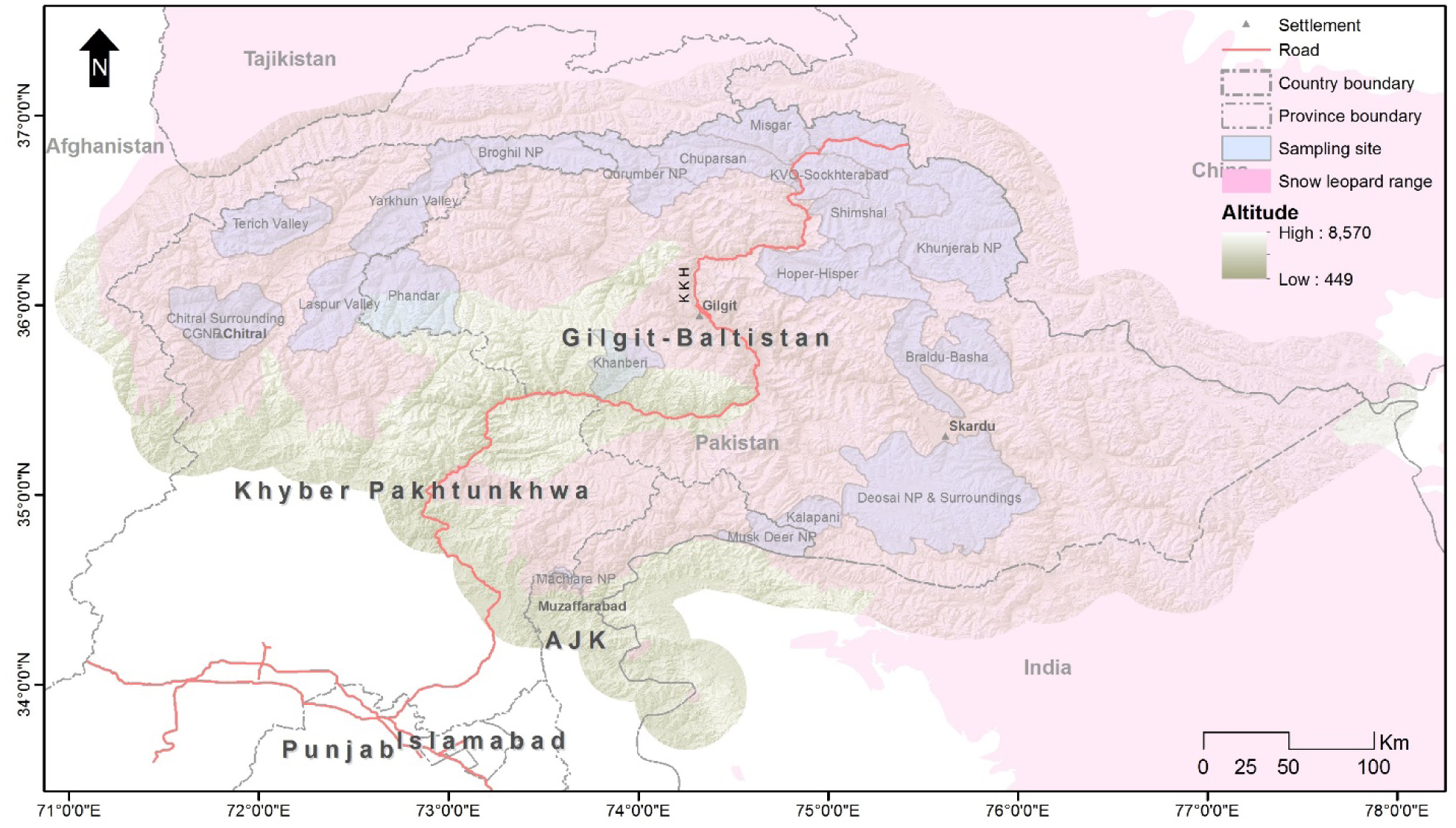
Map of study area showing sampling sites and IUCN range of snow leopard in Pakistan.

**Fig 2.**
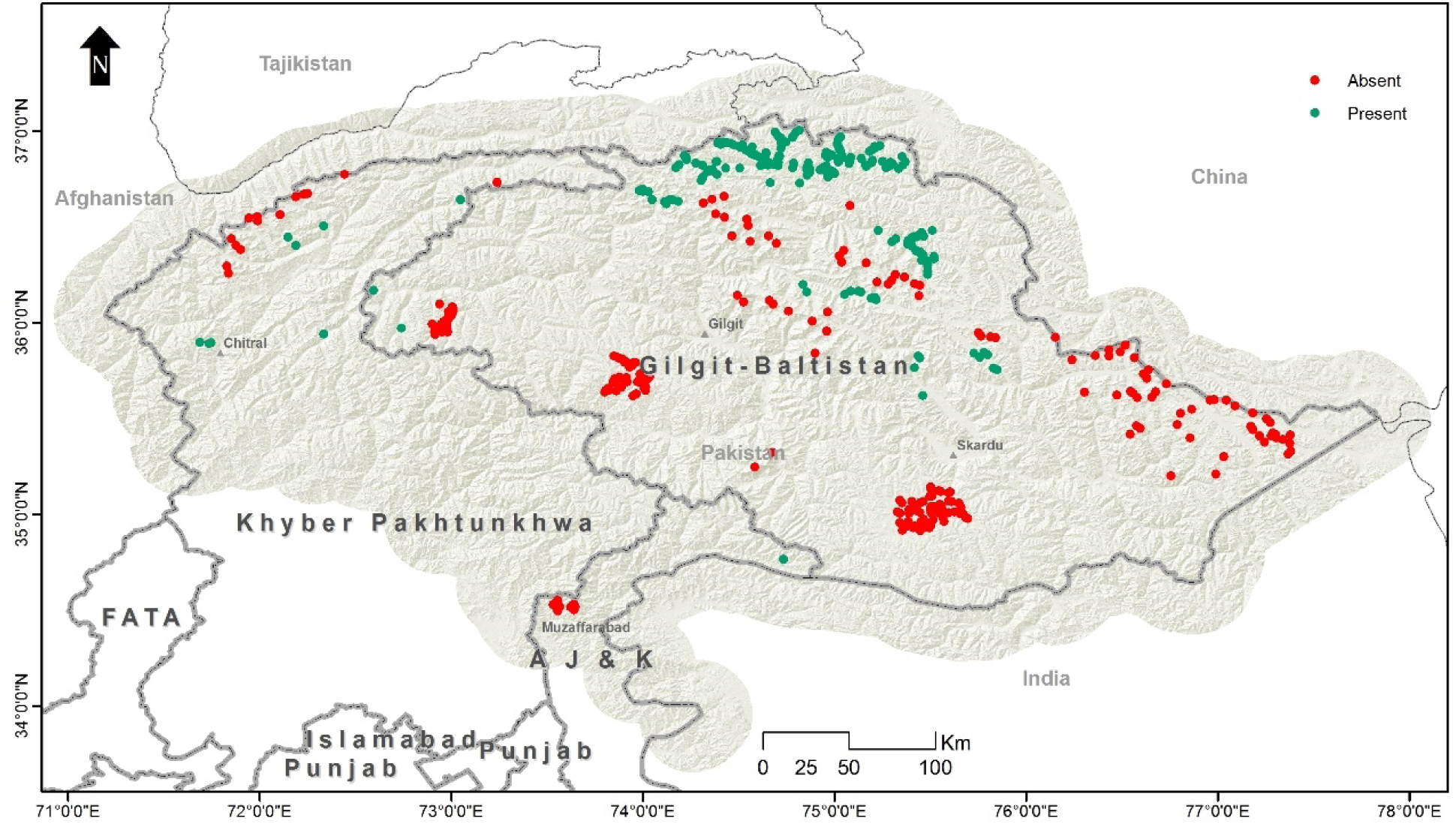
Presence and absence locations of snow leopards used for model evaluation.

These glorious mountain ranges are also home to one of the densest collections of high and precipitous mountain peaks in the world. Their high altitudes and sub-zero temperatures also make them one of the most heavily glaciated parts of the world outside the polar regions. The Western Himalayan Range is situated in AJK and GB to the south and east of the Indus River. The Hindukush rise Southwest of the Pamirs. The Karakoram Range covers the borders between three countries in the regions of GB in Pakistan, Ladakh in India and the Xinjiang region in China. They are considered to extend from the Wakhjir Pass at the junctions of the Pamirs and Karakoram to the Khawak Pass north of Kabul.

The mountainous regions of Pakistan are heavily inhabited despite harsh geographic and climatic conditions. Nevertheless, the special ecological conditions and remoteness of these mountainous areas also support unique biodiversity of plants and animals. Some of these high hills harbour 90% of Pakistan’s natural forests. Climatic conditions vary widely, ranging from the monsoon-influenced moist temperate zone in the western Himalayas to the semi-arid cold deserts of the northern Karakorum and Hindu Kush. Four vegetation zones can be differentiated along the altitudinal ascents, such as alpine dry steppes, subalpine scrub zones, alpine meadows and permanent snowfields. Various rare and endangered animals such as the snow leopard (*Panthera uncia*), grey wolf (*Canis lupus*), brown bear (*Ursus arctos*), Asiatic black bear (*Ursus thibetanus*), Himalayan lynx (*Lynx lynx*), Himalayan Ibex (*Capra ibex sibirica*), blue sheep (*Pseudois nayaur*), flare-horned markhor (*C. f. cashmirensis*), musk deer (*Moschus chrysogaster*), Marco Polo sheep (*Ovis ammon polii*), Ladakh urial (*Ovis orientalis vignei*) Pallas’s cat

(*Otocolobus manual*) and woolly flying squirrel (*Eupetaurus cinereus*), inhabit in these varied climatic conditions and ecosystems.

### Data Collection

Presence records were collected by three methods: camera trapping, sign surveys and genetic sampling. Camera trapping is being increasingly adopted for the monitoring of shy and rare wildlife [35–37]. We deployed 806 camera stations in Chitral Gol National Park (CGNP), the buffer areas of CGNP and Tooshi Game Reserve (TGR), TGR, Laspur Valley, Khunjerab National Park (KNP), Shimshal, Khunjerab Villagers Organization (KVO) area, Qurumber National Park, Broghil National Park, Deosai National Park, Yarkhun Valley, Misgar, Astor, Musk Deer National Park, Khanberi Valley, Terich Valley, Hoper-Hisper, Basha and Arandu and buffer areas of Central Karakoram National Park (CKNP), during the period 2006–2017 (**Fig 1**). These cameras remained active for more than 20,000 trap-days in the field. The camera brands used were CamTrakker™ (Ranger, Wattkinsville, GA, USA) and Reconyx^TM^ (HC500 Hyperfire^TM^ and PC900 Hyperfire^TM^; Reconyx, Holmen, Wisconsin, USA). The sites for camera installation were selected near tracks, scrapes, scats, and other signs. A minimum aerial distance of 1 km was kept between the two nearest camera stations. All essential procedures, safety measures and standards—camera height, front view, sensors, etc.—were followed whilst setting the cameras up, as per Jackson et al. [36]. The majority of the camera stations were supplied with different type of lures—castor, skunk and fish oil—to enhance capture probability.

Site Occupancy based sign surveys were conducted in KNP-KVO-Shimshal, Qurumber- Broghil national parks, Misgar-Chapursan, Phandar Valley, and Basha-Arandu from 2010 to 2017. Each study area was divided into small grids cells of 5 × 5 km—except in KNP-KVO- Shimshal where we kept grid size to 10 × 10 km) on GIS maps. Each grid cell (site) was approached by GPS and multiple points were led to search the signs for snow leopards. A total of 193 sites with 1,607 repeat survey points were searched for signs of snow leopards. Presence was detected through five types of signs (scrapes, pugmarks, faeces, scent spray and claw marks). However, in this analysis, we used just two types of signs, i.e. scrapes and pugmarks, as these are considered more reliable [38].

Faecal samples were collected from 2009 to 2013 during the sign and camera trap surveys. We collected over 1,000 faecal samples of all carnivore species encountered in the field and preserved them in 95% alcohol in 20 ml bottles. The DNA extraction was performed in a laboratory dedicated to the extraction of degraded DNA. Total DNA was extracted from c. 15 mg of faeces using the DNeasy Blood and Tissue Kit (QIAgen GmbH, Hilden, Germany) following the maker’s guidelines with a small modification as explained by Shehzad et al. [39]. Blank extractions were performed to scrutinize contamination. Species identification was performed through next generation sequencings (NGS) by amplifying DNA extract using primer pair 12SV5F (5’-TAGAACAGGCTCCTCTAG-3’) and 12SV5R (5’- TTAGATACCCCACTATGC-3’ targeting about 100-bp of the V5 loop of the mitochondrial 12S gene [37, 39] The sequence analysis and taxon assignation were done using OBITools as described in Shehzad et al. [39, 40].

### Data Analysis

We used MaxEnt modelling [24] to predict snow leopard distribution in Pakistan. MaxEnt is a freely available programme, and we used version 3.3.3k from www.cs.princeton.edu/~schapire/maxent. It predicts species distribution using presence-only data and environmental variables, and estimates species’ probability distribution by finding the probability distribution of maximum entropy, i.e. the most spread out or closest to uniform, subject to a set of constraints [24]. It is amongst the most popular species distribution modelling methods with more than 1,000 published usages since 2005[13, 41]. MaxEnt has also surpassed other methods and exhibited higher predictive accuracy [42].

We used a random seed option and kept 25% data for random tests—25 replicates were run with typeset as a subsample. The rest of the settings were kept as default, which included a maximum of 10,000 background points, 5,000 maximum iterations with a convergence threshold of 0.00001, and a regularization multiplier of 1.

#### Data Preparation

We used snow leopard range with an added buffer of 30 km to model under MaxEnt. All environmental layers were converted to the same size (extent) and resolution, i.e. 1 × 1 km, using ‘resample’, ‘clip’ and ‘mask’ tools in ArcGIS 10.2. Snow leopard occurrence points were also converted into a grid file. All environmental variables and presence points were then converted into ASCII files as required by MaxEnt, by using the ‘conversion’ tool in Arc GIS 10.2. Features in Maxent are derived from two types of environmental variables: continuous and categorical [12].

Among the 28 variables considered initially, 11 environmental variables were retained after a multicollinearity test (Table 2.1), including 4 bioclimatic variables (bio1, bio2, bio12, and bio14), distances from river, roads and settlements, slope, ruggedness, soil and a normalized difference vegetation index (NDVI) [37]. Bioclimatic variables were derived from the mean temperature, minimum temperature, maximum temperature and precipitation in order to generate more biologically meaningful variables—these are often used in ecological niche modelling. Details of each variable used and their sources are shown in **Table 1**.

**Table 1.**
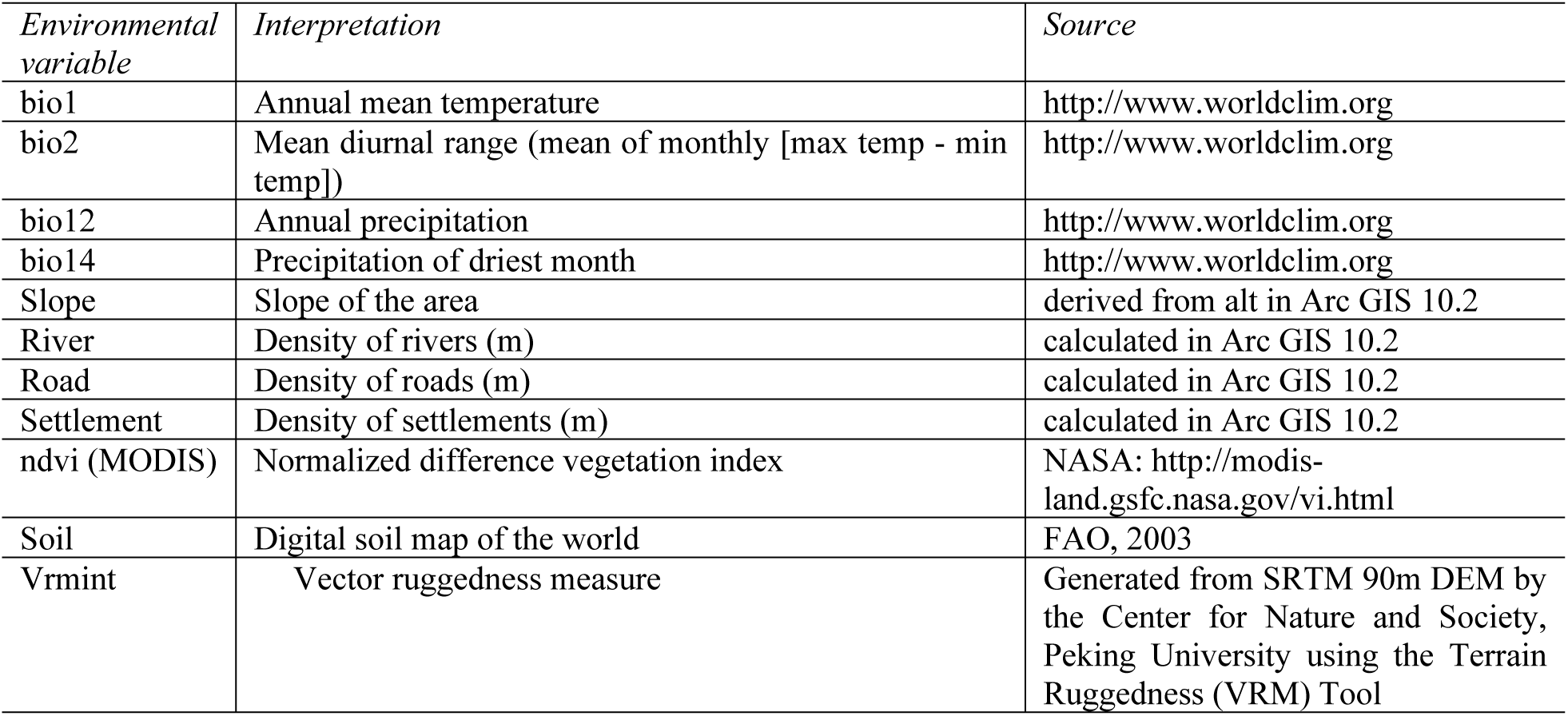
List of environmental variables used in MaxEnt modelling.

#### Model Evaluation

The fit or accuracy of the model should be tested, for every modelling approach, to determine its significance. This can be done in two ways: 1) through receiver operating characteristic (ROC) plots, and 2) defined thresholds [15]. We used both approaches to determine model accuracy.

Model robustness is commonly evaluated by area under the curve (AUC) values of the ROC [43] that range from 0 to 1—AUC values in the range 0.5–0.7 are considered low, 0.7–0.9 moderate, and 0.9, high [44, 45].. Values close to 0.5 indicate a fit no better than that expected by random, while a value of 1.0 indicates a perfect fit. It is also possible to have values less than 0.5—this indicates that a model fits worse than random [46]. It is a graded approach for evaluating model fit that verifies the probability of a presence location being graded higher than a random background locations that serve as pseudo-absences for all analyses in MaxEnt [24]. The AUC quantify the significance of this curve and we used its values to determine model accuracy. ROC is a plot of the sensitivity vs. 1-specificity over the entire range of threshold values between 0 and 1 [47]. Using this method, the commission and omission errors are, therefore, weighted with equal importance for determining model performance [48].

Another approach entails selecting thresholds to determine sites that are considered suitable or unsuitable for the species of interest. These thresholds are established by maximizing sensitivity while minimizing specificity [24]. The proportion of sites that are precisely categorized as suitable locations can be compared to the proportion of unsuitable sites to verify model accuracy. We checked our model output against different defined thresholds and selected the one with the lowest error.

Presence locations excluded by the collinearity model were used for model evaluation along with absence locations. Absence locations were obtained in two ways, a) from surveyed sites where snow leopards were not detected (214 locations), and b) through 102 locations which were extracted from areas higher than 6,500 m—no-go areas for snow leopards [6] (Figure 2.2).

#### Modelling Potential Movement Corridors

Using the snow leopard distribution map generated by MaxEnt, we also modelled for potential movement corridors. This was achieved through Circuitscape 4.0 (software) [49], an open-source programme that uses circuit theory to predict connectivity in heterogeneous landscapes for individual movement. We use pairwise modelling mode which iterates across all pairs in a focal node file. We selected 38 points from different locations and converted them into a grid file in ArcGIS 10.2. Both habitat suitability map (created by MaxEnt) and points files were converted into ASCII format for a Circuitscape model run. We used the option of conductance instead of resistance because, in our model, higher values indicate greater ease of movement and generated cumulative and max current maps, only.

## Results

Snow leopard detection was low as it was photo-captured in 97 capture events at just 60 stations (out of 806 stations) (**Fig 3**). In the majority of our study areas, there was either single capture— Laspur Valley, Qurumber National Park, Musk Deer National Park, Terich Valley—or no capture (Broghil National Park, Deosai National Park, Yarkhun Valley, etc.). Multiple captures occurred only in the Khunjerab National Park, Shimshal, and Misgar valleys, Hoper-Hisper, and buffer areas of Central Karakoram National Park.

**Fig 3.**
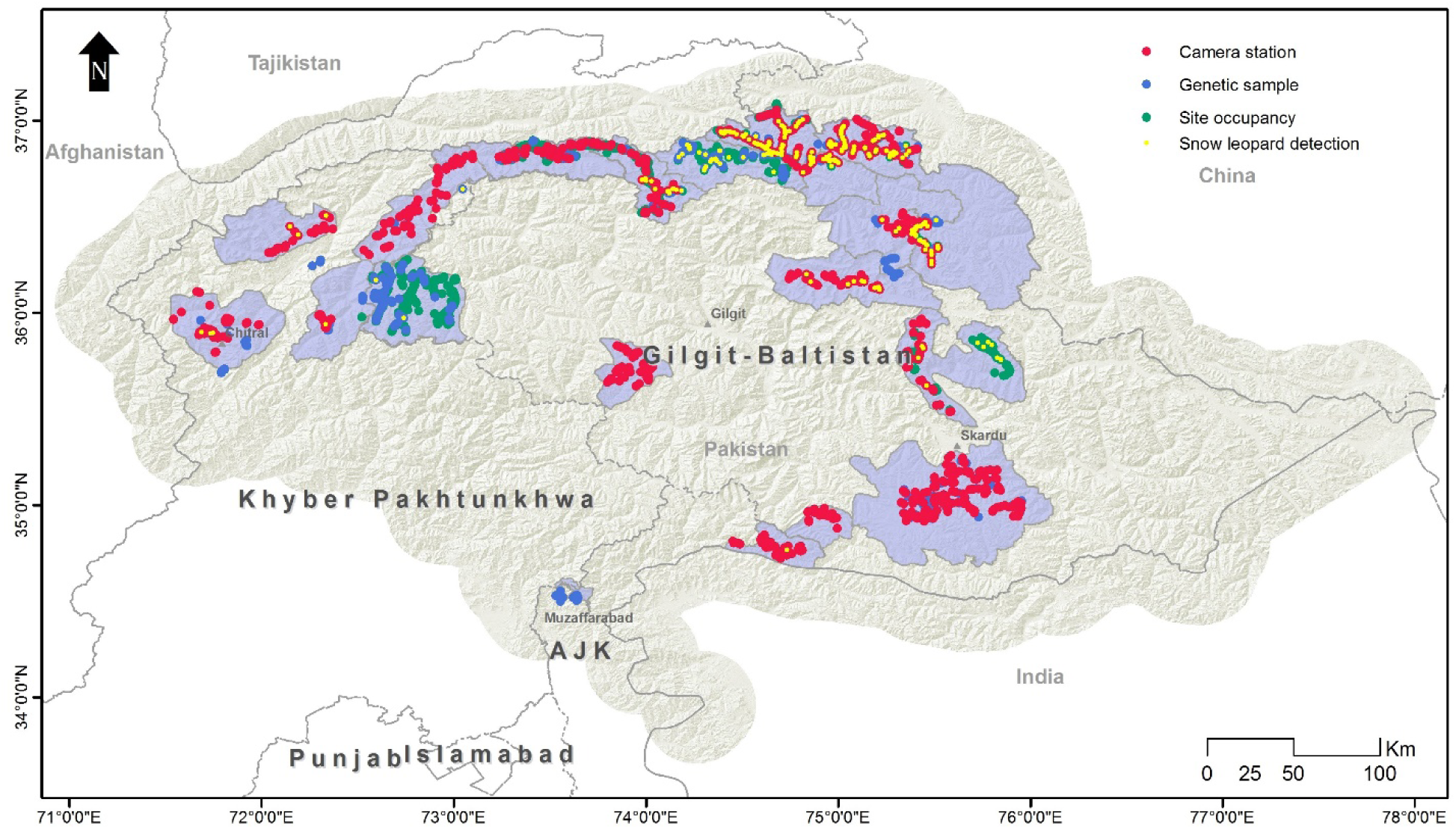
Sampling locations by different survey techniques and snow leopard detections.

In sign-based site-occupancy surveys, signs older than ten days were also excluded to avoid misperception. After this screening, we obtained 213 locations in different areas with fresh signs—either scraped or pugmarks, or both. Among 1,000 faecal samples, a genetic analysis confirmed 111 to be of snow leopards. Combining all three methods, we obtained 384 (Figure 3.1) confirmed locations of snow leopards. These locations were overlapping in some areas where multiple surveys were conducted. Records obtained by signs, scats and camera trappings were screened in SDMtoolbox, a tool of GIS, to remove spatially correlated data points to guarantee independence [50–52] After this selection, 98 unrelated locations were used to generate current SDMs of the snow leopard.

### Range-wide Habitat Suitability

MaxEnt produced outputs for 25 replicates and averaged them into one model along with response curves and AUC. This average model was used for interpreting habitat suitability and calculating potential movement corridors.

The habitat suitability score calculated by MaxEnt ranged from 0 to 0.97 across the snow leopard’s assumed range in Pakistan (**Fig 4**). A large shear of previously known range represented low-quality habitat, including areas in lower Chitral, Swat, Astor and AJK. Conversely, KNP, Misgar, Chapursan, Qurumber National Park, Broghil National Park, and CKNP represented high-quality habitats.

**Fig 4.**
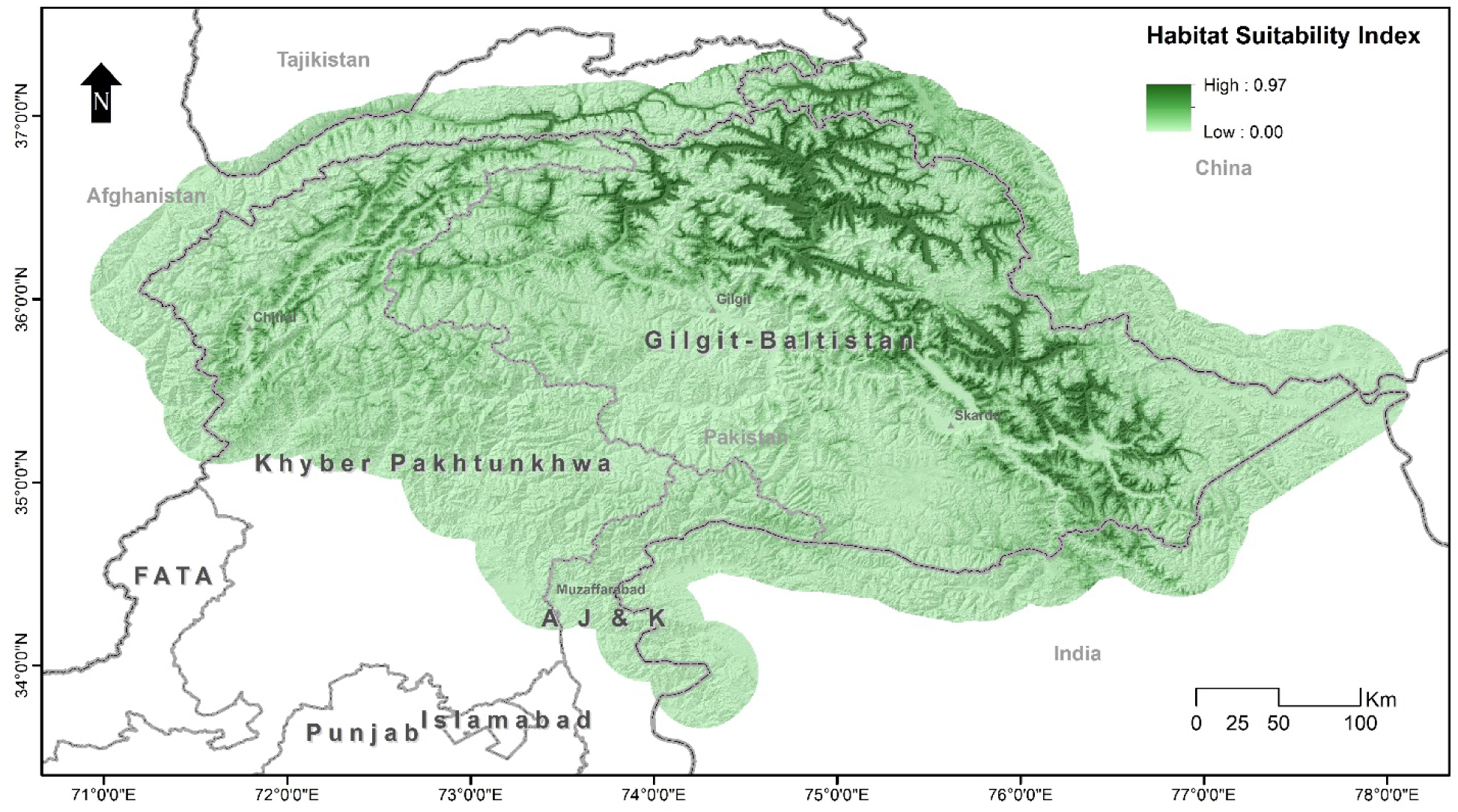
Habitat suitability of snow leopards in Pakistan, calculated with MaxEnt.

**Fig 5:**
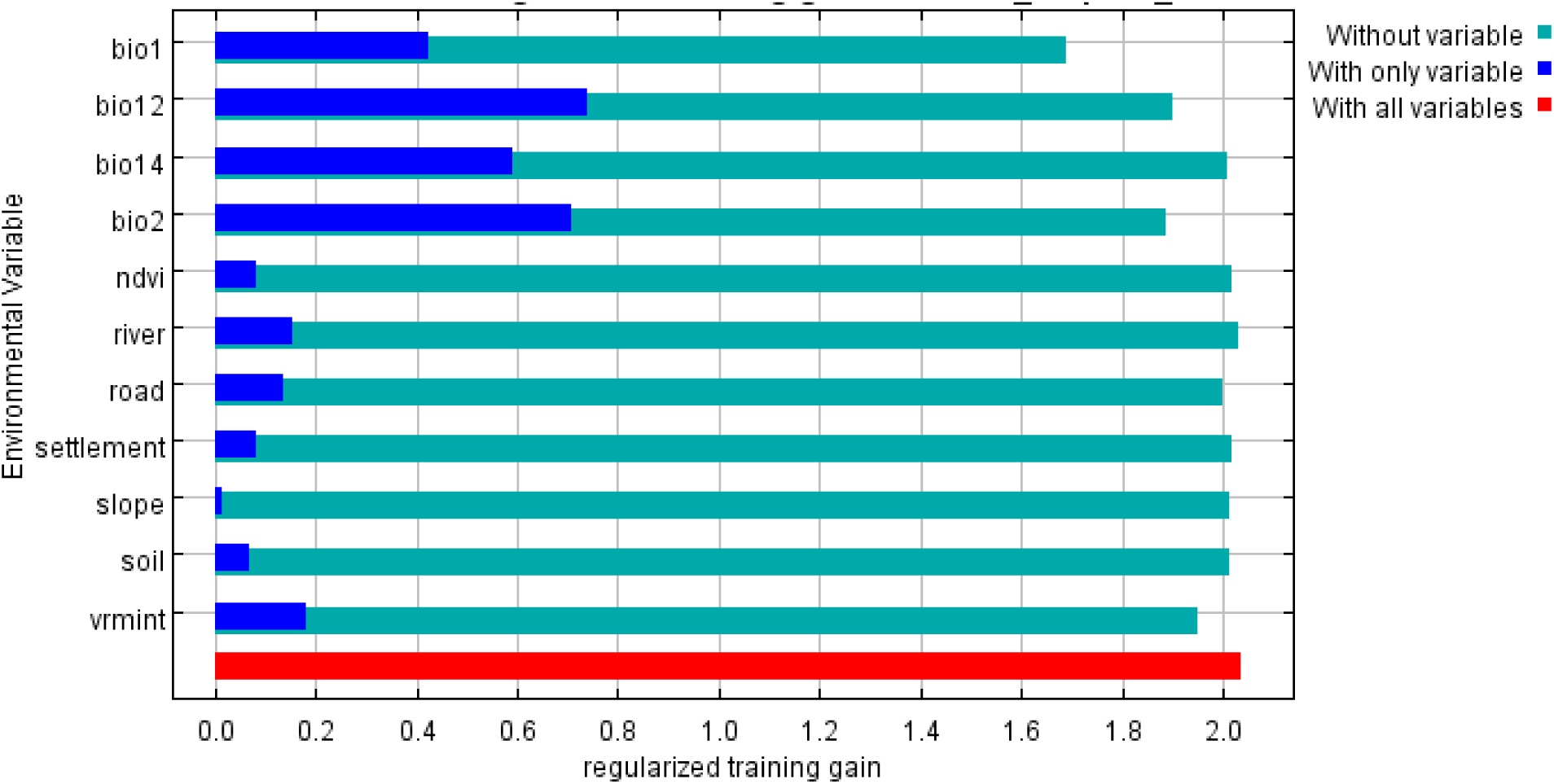
Jackknife test of regularized training gain of variables tested in snow leopard habitat suitability model.

### Factors Determining Habitat Suitability

Variables with higher contribution in the MaxEnt model were precipitation of driest month (34%), annual mean temperature (19.5%), mean diurnal range of temperature (9.8%), annual precipitation (9.4%) and river density (9.2). The contribution of other variables included in the model was low (**Table 2**).

**Table 2.**
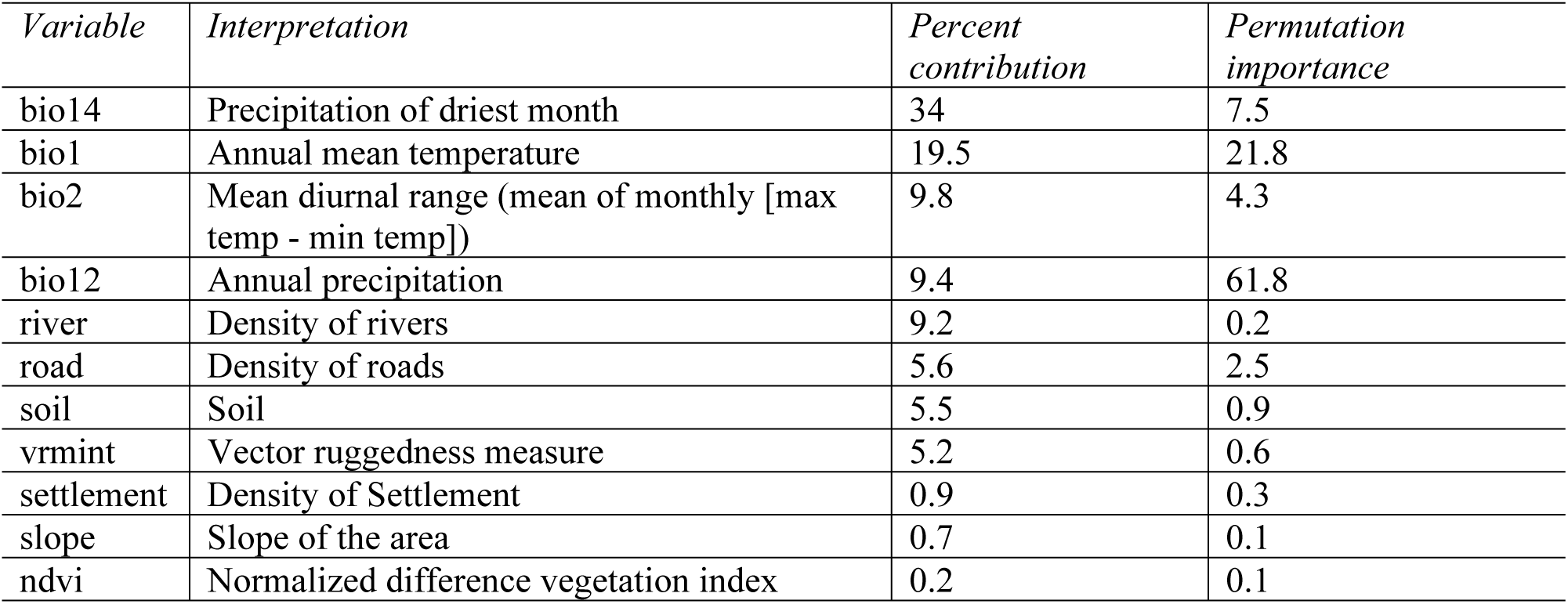
Estimates of relative contributions of the environmental variables to the Maxent model.

The Jackknife Test of variable importance showed that the environmental variable with the highest gain when used in isolation is density of river, which, therefore, appears to have the most useful information by itself. The environmental variable that decreased the gain the most when it was omitted was annual mean temperature (bio1), which, therefore, appears to have the most information that is not present in other variables. The values shown are averages over replicate runs. (**Fig 4**).

### Model Evaluation and Threshold Selection

MaxEnt performed some basic statistics on the model and calculated an averaged AUC for the model. Analysis of omission/commission was done by MaxEnt and Figure 3.6a shows the test omission rate and predicted area as a function of the cumulative threshold averaged over the replicate runs. The omission rate should be close to the predicted omission because of the definition of the cumulative threshold and, in our case, is very close to the predicted one.

The ROC curve (**Fig 6b**) for the data was also calculated by MaxEnt, again, averaged over the replicate runs. Here, specificity is defined using predicted area rather than true commission [24]The average test AUC for the replicate runs was 0.933 and the standard deviation was 0.024.

**Fig 6.**
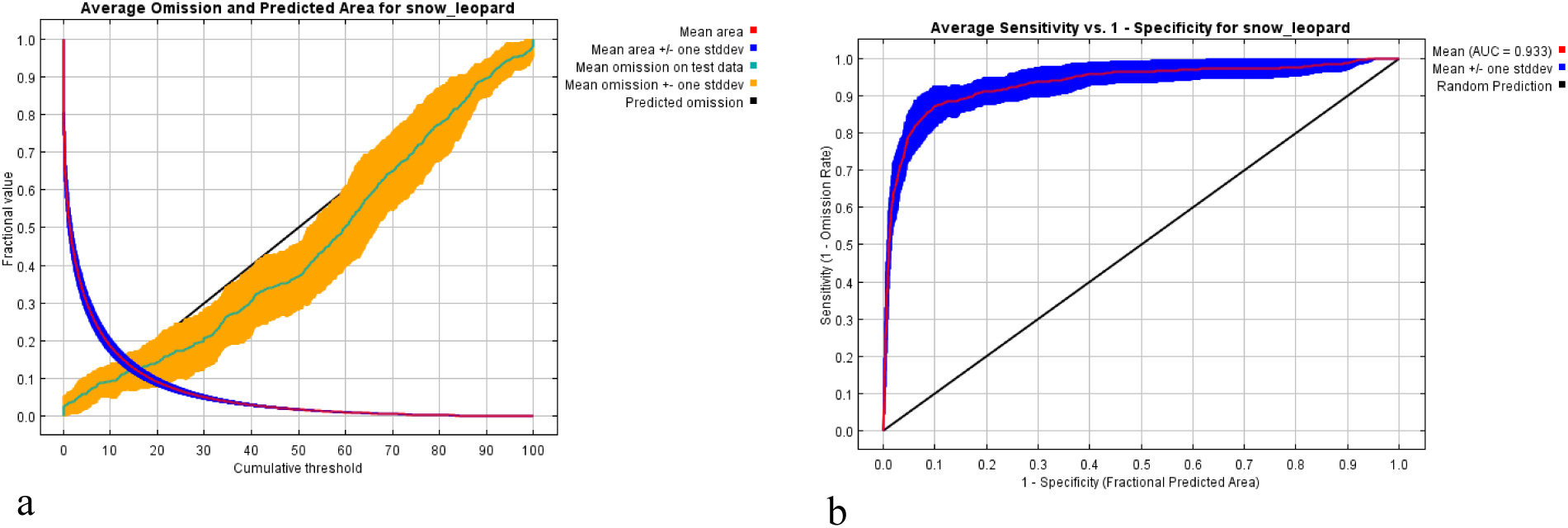
Model evaluations, (a)Averaged omission and predicted area for snow leopard, (b) The ROC curve calculated by MaxEnt as averaged sensitivity versus 1-specificity for snow leopard.

**Fig 7.**
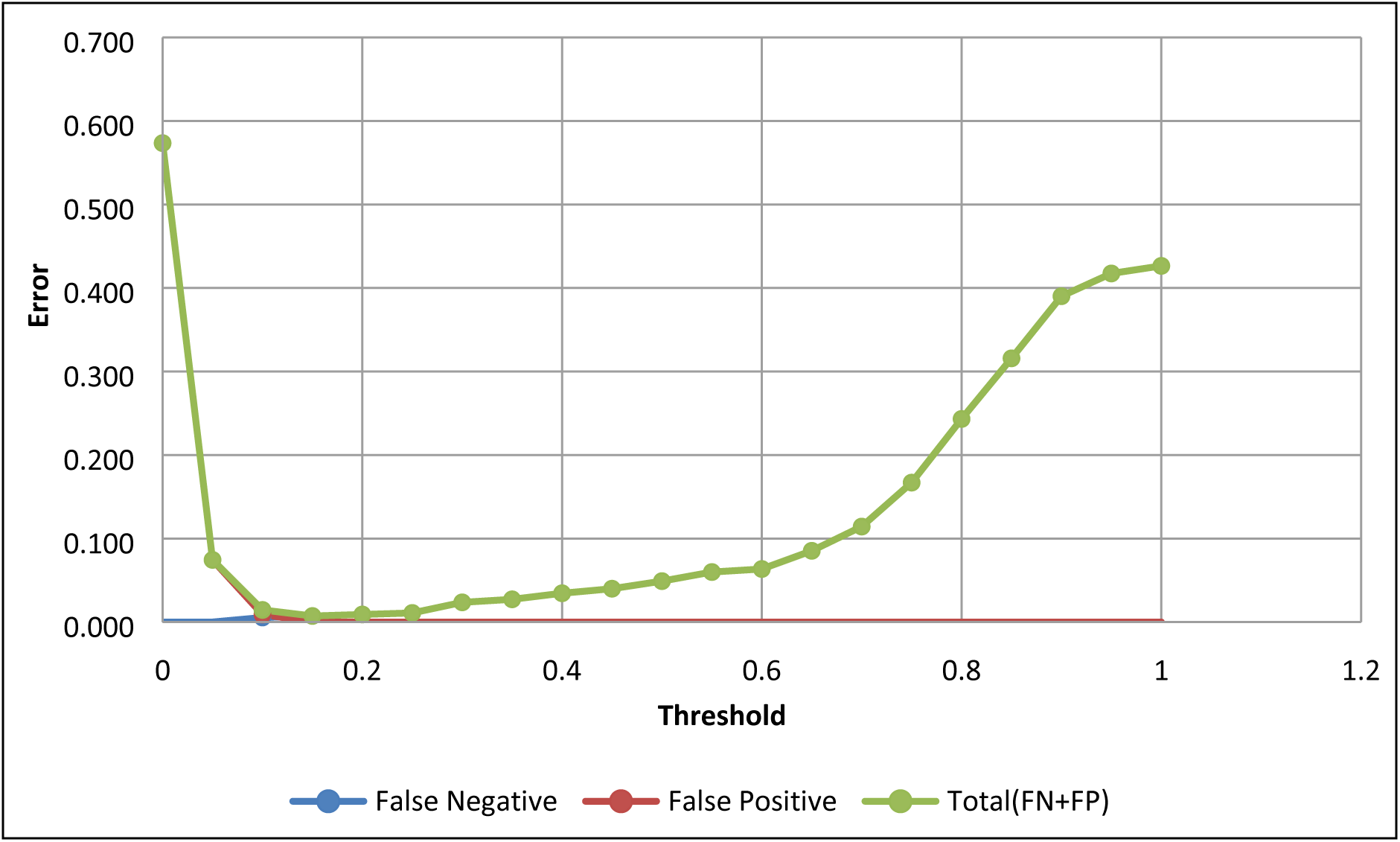
Graph showing the relationship of False Negative and False Positive rates against different thresholds of model prediction.

Measuring the error of false positive (FP) and false negative (FN) rates against a range of defined thresholds (Figure 3.8), the lowest error was found at a threshold of 0.15. The binomial map was re-evaluated by plotting presence and absence points and it showed that almost all presence points were in suitable habitat areas and absence points in unsuitable areas. The values of 235 presence points and 316 absence points were extracted from the model and plotted against different thresholds. The value of AUC by ROC curve calculated at 0.15 was 1.000; which means our model performed very well.

It was calculated that 235 points were true positives (TPs) and 275 were true negatives (TNs), while FPs were 41 and FNs were 0. The true positive rate (TPR) was calculated at 1.000 while the false positive rate (FPR) was 0.130. Accuracy and specificity were calculated at 0.926 and 0.870, respectively, while the positive predictive value (PPV) was found to be 0.851 and the negative predictive value (NPV) was 1.000. The false discovery rate (FDR) was calculated at 0.149.

### Potential Movement Corridors of the Snow Leopard

The circuit model (**Fig 8**) revealed an interesting pattern with respect to the snow leopard’s habitat connectivity. The population in the Hindukush landscape appears to be more connected with the population in Afghanistan as compared to other populations in Pakistan. Similarly, the Pamir-Karakoram population is better connected with China and Tajikistan, and the Himalayan population with the population in India.

**Fig 8.**
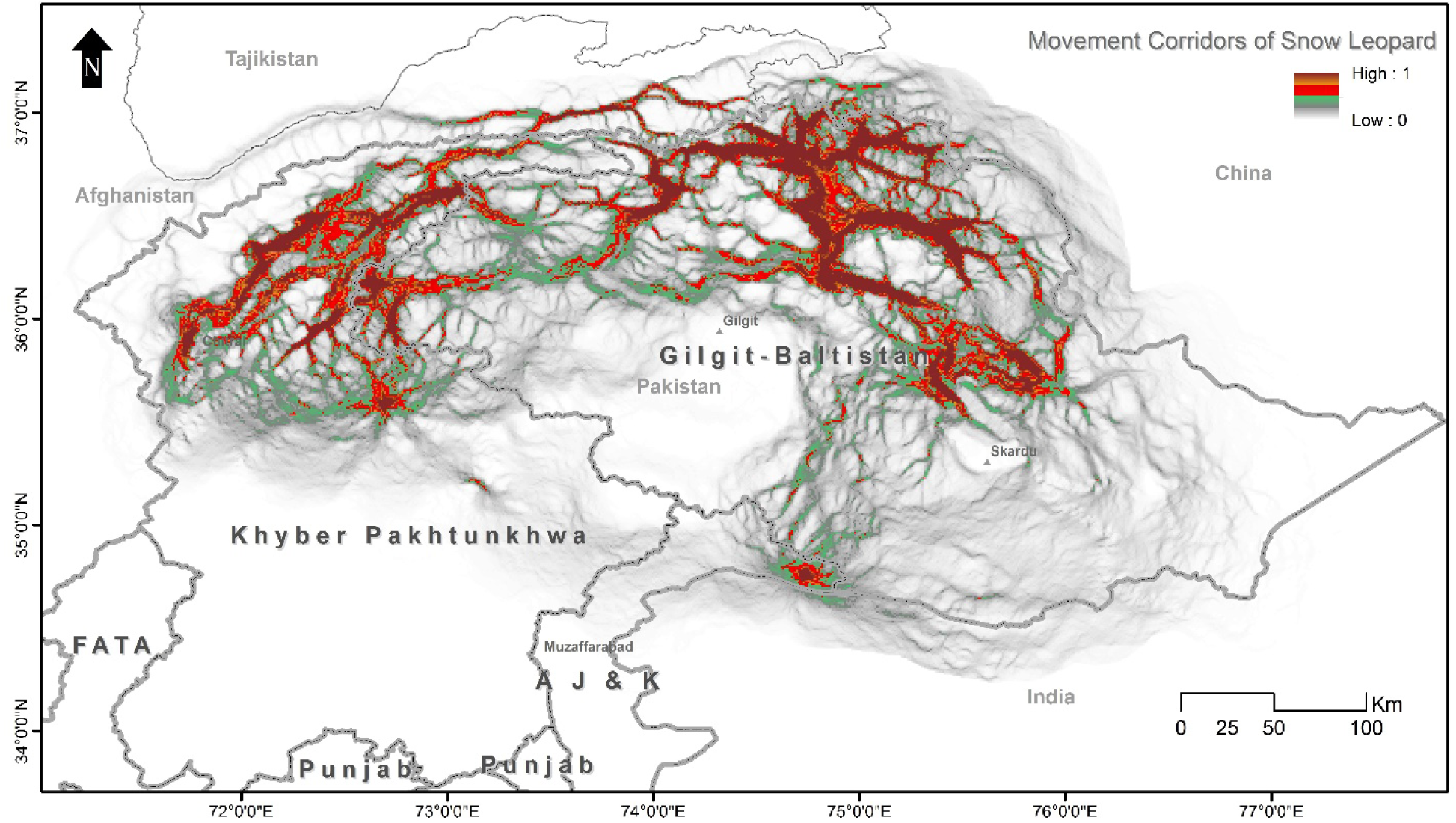
Potential movement corridors of snow leopards in northern Pakistan calculated through Circuitscape.

We observed that Chitral had weak connections with other areas when we examined habitat connectivity in Pakistan. However, the populations of Phandar, Laspur Valley and Yarkhun Valley seemed connected. Interestingly, Broghil National Park had a weak connection with its adjacent Qurumber National Park, but had strong links with Yarkhun Valley, while Qurumber National Park had strong links with Chapursan which is connected to Misgar, which had a strong link with KNP. The populations of CKNP and Musk Deer National Park were also shown to be isolated from others and the latter did not even have any movement corridors close to it.

### Protected Areas Coverage in Snow Leopard’s Habitat in Pakistan

Habitat Suitability model was also assessed against current protected area coverage (**Fig 9**). It was revealed that most of the the suitable habitat of snow leopard in Paksitan has already been protected, however there are some areas like Misgar, Chipursan and Terich that are outside of any declared protected area.

**Fig 9.**
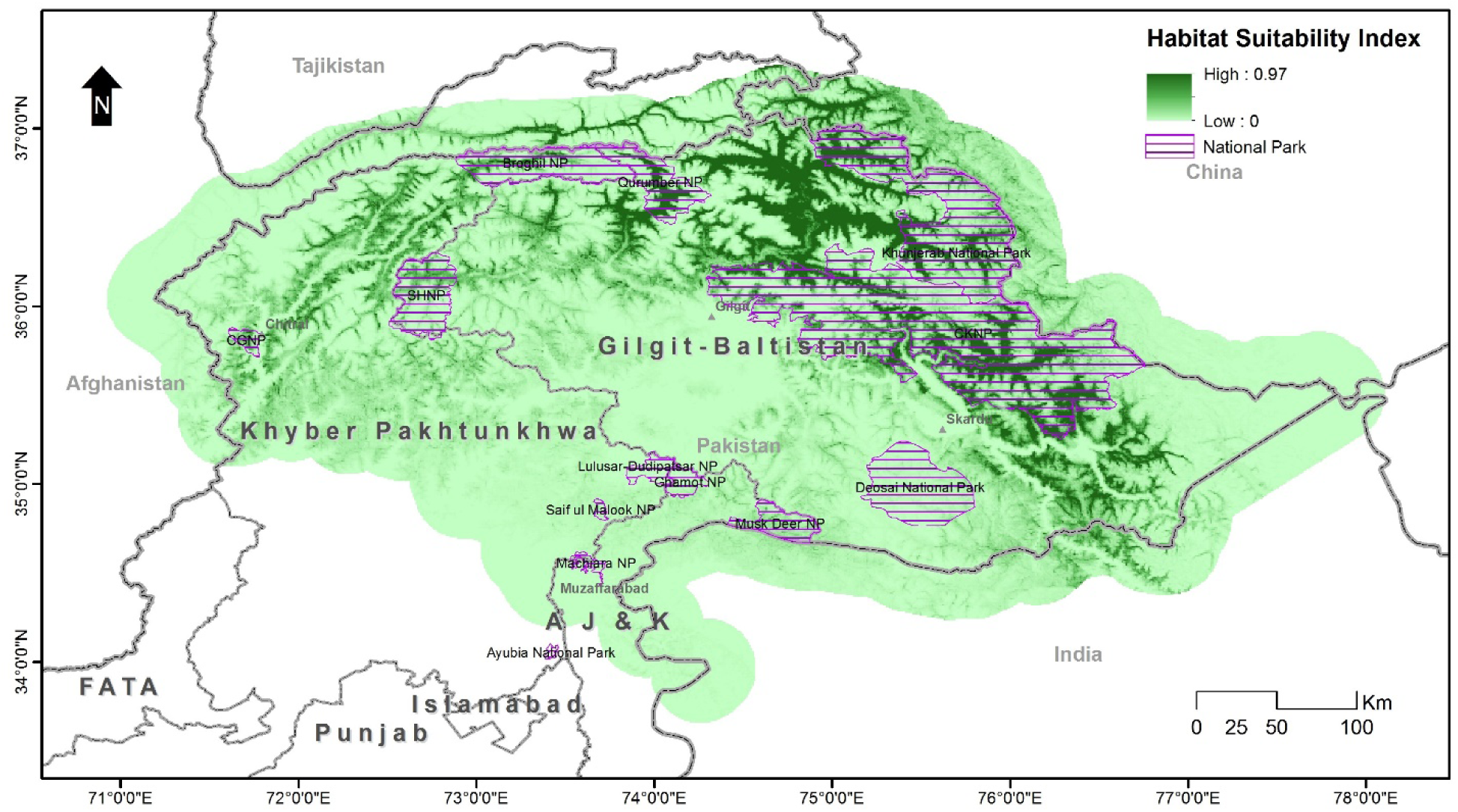
Snow leopard habitat versus protected area coverage.

It was also observed that most of the national parks had weak links with regards to movement of snow leopard across different habitats (**Fig 10**). Even some adjascent protected areas, like; Broghil-Qurumber National Parks and Khujerab-Central Karakoram National Parks had no or very weak movement corridors of snow leopard at their shared borders.

**Figure 10.**
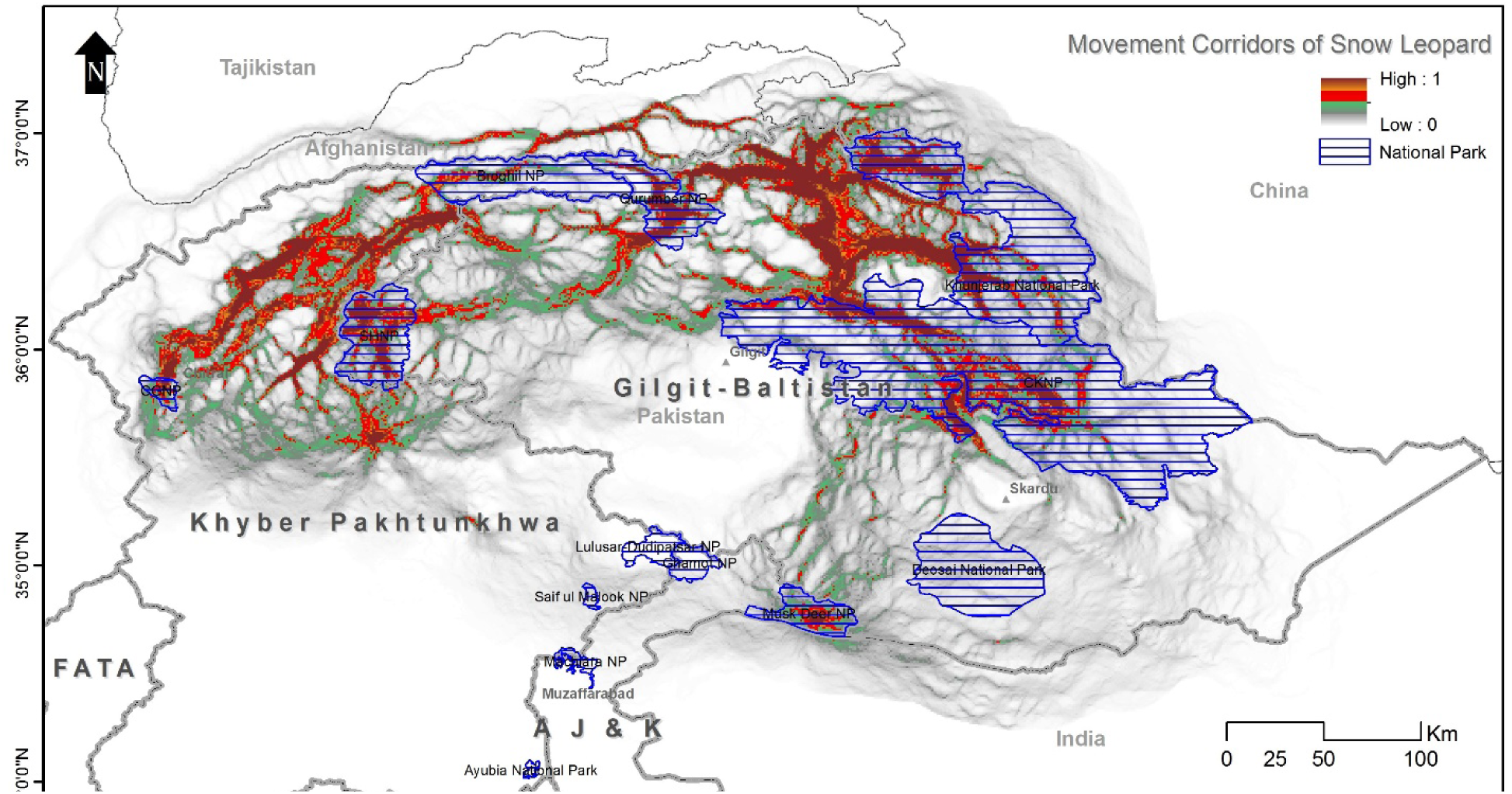
Movement corridors between different National Parks.

## Discussion

This is the first known study on the snow leopard’s distributional patterns and habitat connectivity in Pakistan—it revealed some interesting facts about the species’ habitat in Pakistan. It was observed that the cat’s distributional range mentioned in Roberts [10] and Fox [11] had very weak scientific grounds, which was obvious due to the lack of data available at the time. We recorded snow leopard presence using multiple techniques and discarded all ambiguous entries. Moreover, we surveyed over 25,000 km^2^ which covered about 30% of known snow leopard range in Pakistan. In addition, we did not limit our surveys and model to just snow leopard range but extended it to a 30% buffer area to include potential areas that could possibly be favoured by snow leopards for their movement. The study showed that snow leopard presence is not restricted to its known range and that it possibly uses other areas as well. We discovered that numerous areas in the snow leopard range either have very low suitability or are unsuitable for its presence.

Although our dataset was vast, we used presence records only to predict the habitat suitability model as it has the advantage of being derived from different sources that can be combined to inform control projects [27]. This released us from the problems of unreliable absence records [38]. Modelling applications like Maxent [24] are highly suitable for predicting species’ distribution based on available presence records without model under-fitting [15, 53]. This model, which is one of the most widely used ones to model species distributions, is a machine-learning method based on maximum entropy. Absence data is replaced with ‘background data’ or ‘pseudo-absences’ which are a random sample of the available environment. Maxent estimates a target probability distribution by finding the probability distribution of maximum entropy and its logistic output can be used as a habitat suitability index [12, 54].

We selected Maxent because it typically outperforms other methods based on predictive accuracy and the software is particularly easy to use [41, 55]. Since becoming available in 2004, it has been utilized extensively to model species distributions [38]. Several studies were undertaken to compare the results of Maxent with other methods and it was found that Maxent predicted suitable areas better than regularized logistic regressions based on the expert-based landscape classification [27]. Maxent was also used to predict the distribution of snow leopard in various countries [52, 56].

This study showed that most of the snow leopard’s habitat is patchy, having no or weak links with other areas. Though, there are potential movement corridors between different areas, e.g., between KNP and CKNP, but these are not strong enough to be called permanent routes (Figure 3.12). The connectivity model also revealed that in some areas, snow leopard possibly favoured movement across borders instead of inside Pakistan, e.g., Broghil National Park had more connectivity to Afghanistan than to its adjacent national park, Qurumber National Park. Also, KNP and CKNP did not show any connectivity at their shared border, but there is a movement corridor between these two parks on the other side. These connectivity patterns seem unusual on maps, but other factors like the presence of large glaciers explain the absence of any movement corridors at the borders of these parks. This connectivity model proposed by McRae et al. [32] from electrical circuit theory is a useful addition to the approaches available to ecologists and conservation planners. Circuit theory can be applied to predict the movement patterns and probabilities of successful dispersal or mortality of random walkers moving across complex landscapes, to generate measures of connectivity or isolation of habitat patches, populations, or protected areas, and to identify important connective elements (e.g., corridors) for conservation planning [32]. The establishment of movement corridors can offset the negative effects of habitat fragmentation by connecting isolated habitat populations or patches [57, 58].

Our habitat suitability model was also useful for assessing the coverage of protected areas, specifically national parks in the snow leopard’s habitat. Although a lot of suitable snow leopard area falls in national parks, there are still many areas that need to be included in the protectead areas network (**Fig 9**), in order to safeguard longetm future of the species. Misgar and Chapursan falling between KNP and Qurumber National Park are some of the most suitable areas for snow leopards that need protection. Areas on the eastern side of CKNP are also not protected. Qurumber National Park is unique in the sense that its entire area is favourable for snow leopards. But there should be a new protected area or extension of Qurumber National Park on its southern and southwest side. Yasin Valley is another important area adjacent to the southern side of Broghil National Park that requires protection. The upper part of Chitral district in KP province is also suitable for snow leopards yet in need of protection.

## Recommendations

The Global Snow Leopard Ecosystem Protection Program (GSLEP) is joint initiave of 12 snow leopard range countries, established to safeguard snow leopards and their the vast ecosystem. The overall aim of GSLEP is to secure at least 20 snow leopard landscapes (SLL) across the cat’s range [33]. Among these 20 model landscapes, three were prposed in Pakistan. Each SLL is defined as an area that can support at least 100 snow leopards of breeding age, has adequate and stable prey populations, and has functional connectivity to other snow leopard landscapes, including across international boundaries [33]. However, in reality, the definition of these landscapes are theoretical, and their boundaries are marked on limited information except for few areas where empirical adat were available. Current study allows us to propose three model landscapes to be included in the GSLEP agenda, based on habitat suitability of the snow leopards across Pakistan. There are named after mountain ranges they fall in; Himalaya, Karakoram- Pamir and Hindukush (**Fig 11**). We also recommend Government of Pakistan to establish new national parks to protect critical habitats of snow leopards falling in Misgar, Chuparson and Terichmir areas in Gilgit-Baltistan and Chitral.

**Fig 11.**
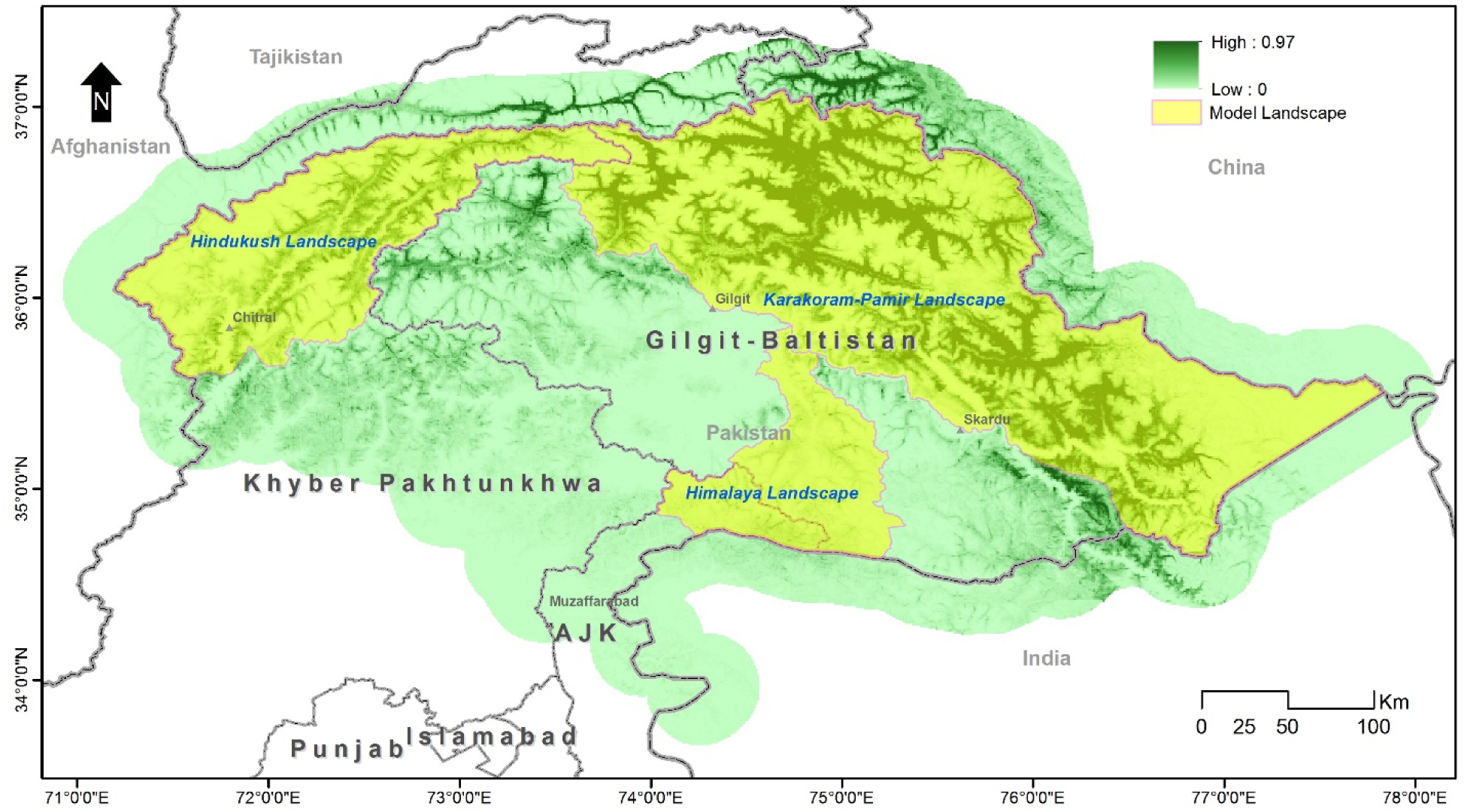
Recommended model landscapes for GSLEP.

## Acknowledgments

The logistics support provided by provincial wildlife departments of Gilgit-Baltistan, Khyber Pakhtunkhwa, and Azad Jammu and Kashmir, and local communities is highly appreciable. We acknowledge the support of Snow Leopard Foundation for field work in terms of equipment and trained field staff. We are highly grateful to Muhammad Ayoub, Khurshid Ali Shah, and Siraj Khan to help in field work specially camera trapping and sign surveys). We are thankful to Doost Ali and Center for Nature and Society, Peking University for providing much needed GIS support and data. Shakeel Ahmad helped in formatting and editing.

